# Chronic stress does not induce behavioural signs of tinnitus or cochlear synaptopathy in Mongolian gerbils

**DOI:** 10.64898/2026.07.22.739982

**Authors:** David Hein, Konstantin Tziridis, Jwan Rasheed, Claudia Böhm, Holger Schulze

## Abstract

Tinnitus is commonly associated with psychosocial stress. However, it is unclear whether stress alone is sufficient to induce neural alterations resulting in tinnitus. In this study, we investigated the causal role of stress in tinnitus generation by experimentally isolating stress exposure as the sole independent variable in a well-established Mongolian gerbil model. Animals were subjected to repeated, inescapable electric foot shocks over three weeks to create a chronic, repeated stress paradigm. Behavioural, endocrine and histological measures were combined to assess tinnitus perception, stress system activation and cochlear synaptopathy.

Repeated stress exposure reliably activated the endocrine stress response, as indicated by transient increases in serum cortisol following specific stress sessions. Basal cortisol levels returned to baseline values after the stress period. Behavioural assessment using gap-prepulse inhibition of the acoustic startle reflex revealed only isolated cases of behavioural signs of tinnitus, which occurred at a frequency consistent with false-positive detection and were evenly distributed between stress-exposed and control animals. Histological analyses revealed that ribbon synapse counts were preserved across the cochlea, with no evidence of stress-induced synaptopathy. No systematic relationships were observed between endocrine activation, behavioural outcomes and the number of inner hair cell ribbon synapses.

Taken together, these findings suggest that, in the absence of acoustic trauma, repeated chronic stress is not sufficient to induce tinnitus or inner hair cell synaptopathy. These results argue against a primary causal role of stress alone in tinnitus generation and instead support models in which stress modulates symptom expression or perceptual salience in the context of pre-existing auditory dysfunction.

## 1 INTRODUCTION

Tinnitus, defined as the conscious perception of sound in the absence of an external acoustic source, affects approximately 12–30% of adults and constitutes a substantial clinical and socioeconomic burden (McCormack *et al*., 2016; Stockdale *et al*., 2017; Tziridis *et al*., 2022). Although often transient, a significant subset of patients develops chronic and distressing tinnitus accompanied by sleep disruption, cognitive impairment, and affective comorbidities, including anxiety and depression (Langguth *et al*., 2013; Pattyn *et al*., 2016). Despite its prevalence and impact, the neurobiological mechanisms that initiate tinnitus remain incompletely understood.

Contemporary models conceptualize tinnitus as a disorder of maladaptive auditory plasticity. Reduced or distorted peripheral input is thought to drive compensatory increases in central neural gain and neuronal synchronicity, trigger homeostatic plasticity or stochastic resonance, disrupt inhibition, drive large-scale network reorganization, and/or generate prediction errors, ultimately generating persistent phantom percepts (Eggermont & Roberts, 2015; Knipper *et al*., 2020; Knipper *et al*., 2021; Schilling *et al*., 2023; Schulze & Schilling, 2026). Cochlear synaptopathy - the loss of ribbon synapses between inner hair cells and auditory nerve fibers - has emerged as a critical peripheral trigger capable of inducing such central adaptations (Liberman & Kujawa, 2017). Consistent with this framework, animal models based on acoustic trauma or ototoxic damage reliably produce both synaptic deafferentation and tinnitus-like behavioural phenotypes, supporting a causal link between peripheral injury and central maladaptation (Rüttiger *et al*., 2013; Tziridis *et al*., 2021).

Beyond peripheral auditory pathology, psychosocial stress has been increasingly discussed as a contributing factor to tinnitus vulnerability and chronicity. Individuals with chronic tinnitus consistently report higher perceived stress levels and show elevated rates of anxiety and depressive symptoms relative to non-tinnitus controls, indicating a close association between tinnitus burden and affective distress (Hébert & Lupien, 2007; Trevis *et al*., 2016). These clinical findings are paralleled by alterations in physiological stress regulation, including atypical hypothalamic-pituitary-adrenal (HPA) axis activity and altered cortisol secretion profiles at baseline or in response to acute stressors (Hébert *et al*., 2004). At the neurobiological level, stress activates the HPA axis, leading to systemic glucocorticoid release (Sapolsky *et al*., 2000). Glucocorticoid receptors are expressed in both central auditory structures and the cochlea, providing anatomical substrates through which circulating stress hormones may influence auditory processing (McEwen, 1998; Canlon *et al*., 2007; Tahera *et al*., 2007). Chronic elevations in circulating cortisol may therefore influence cochlear function and potentially increase vulnerability to stress-related dysregulation and contribute to maladaptive auditory plasticity, highlighting the importance of assessing cochlear integrity to distinguish central from peripheral contributions. Consistent with this interaction, stress-related biomarkers such as cortisol have been associated with the severity and distress of tinnitus, suggesting a relationship between neuroendocrine stress regulation and symptom expression (Mazurek *et al*., 2012). Despite this clinical and biological convergence, existing evidence remains largely correlational and whether stress directly contributes to tinnitus generation or instead represents a secondary consequence of chronic symptoms remains one of the most important unresolved questions in tinnitus research.

Animal models provide a tractable framework to test this causal relationship. The Mongolian gerbil (Meriones unguiculatus) offers a human-like hearing range and well-characterized auditory physiology (Ryan, 1976; Gaese *et al*., 2009) and tinnitus-like perception can be quantified behaviourally using gap prepulse inhibition of the acoustic startle reflex (GPIAS) (Turner *et al*., 2006; Schilling *et al*., 2017). Combining behavioural readouts with endocrine markers and histological quantification of ribbon synapses allows direct linkage of stress signalling to both functional and structural correlates of auditory deafferentation.

We experimentally isolated stress exposure in the absence of acoustic trauma to determine whether stress alone is sufficient to induce tinnitus-like behaviour and cochlear synaptopathy. By integrating behavioural (GPIAS), endocrine (circulating cortisol) and anatomical (ribbon synapse counts) measurements, we systematically assessed the impact of neuroendocrine stress signalling across functional and structural levels of the auditory system. This design enables a direct test of the causal role of stress in tinnitus initiation and distinguishes between models proposing stress as a primary driver of tinnitus versus those in which stress primarily modulates symptom severity or distress.

## 2 MATERIALS & METHODS

### 2.1 Ethical statement

A total of 23 male Mongolian gerbils (Meriones unguiculatus) aged 8–12 weeks were obtained from Janvier (Saint Berthevin, France) and were housed in groups of 2–4 animals per cage in type IV cages within a ventilated housing system (UniProtect Air Flow Cabinet, Zoonlab, Castrop-Rauxel, Germany) under controlled environmental conditions and a 12/12-h light/dark cycle. Animals received standardized commercial pelleted diet for gerbils and water ad libitum. Environmental enrichment was provided in the form of nesting material and additional structural elements. Animals were regularly weighed and closely monitored throughout the study. Three animals were excluded prior to experimental testing due to spontaneous seizure activity. Before the start of the experiments, animals were routinely handled and habituated to the experimental procedure and testing. All experimental procedures involving animals were approved by the State of Bavaria (Regierungspräsidium Unterfranken, Würzburg, Germany; approval no. 55.2.2-2532-2-1350).

### 2.2 Experimental design and procedure

Animals were randomly assigned to one of two experimental groups, stress and control, respectively; due to the nature of the intervention, blinding was not feasible. Following the exclusion of three animals prior to the onset of experiments, each group comprised 10 animals.

The experimental procedure was as follows (cf. Figure 1): Before the stress paradigm was applied, baseline measurements were carried out on all animals. These included GPIAS measurements for subsequent tinnitus assessment, auditory brainstem response (ABR) recordings to determine hearing thresholds and blood sampling for quantitative cortisol analysis.

**Figure 1.**
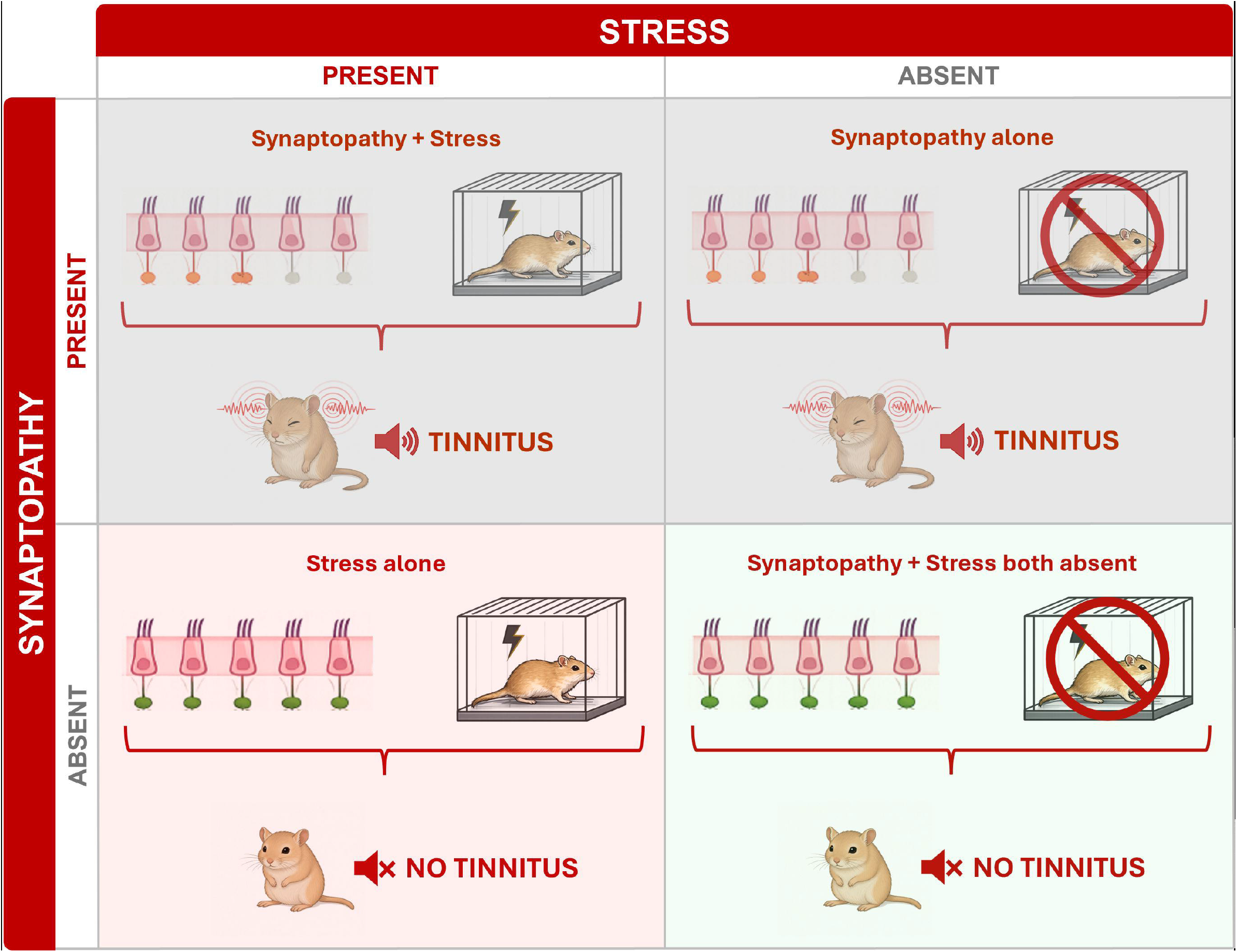
Experimental design and timeline. Animals underwent baseline auditory, behavioural and endocrine assessments prior to the experimental intervention. Stress-exposed animals were subjected to repeated inescapable electric foot-shock stress (IFS; 5x/week) over three weeks. Blood samples (BS) for cortisol analysis were collected at defined time points during the stress period (after day 1, day 5, day 10, and day 15 of IFS). BS, GPIAS and ABR were repeated after the stress period, followed by CO_2_euthanasia and cochlear tissue collection for immunohistological analysis (IH). Control animals underwent identical handling without stress exposure.

Animals from both groups were then exposed to a series of five consecutive daily sessions within a dedicated experimental cage for a period of three weeks. During the first session, animals assigned to the stress group received an extended stimulation protocol (30 electric shocks over 15 min), whereas all subsequent sessions consisted of 10 shocks over 10 min. Control animals underwent identical experimental conditions and duration, with no electrical stimulation applied. Blood samples for cortisol determination were collected from all animals after the 1st, 5th, 10th, and 15th stress session.

At the commencement of the fourth week, blood sampling, ABR measurements for hearing threshold assessment and follow-up GPIAS recordings were repeated. Following the CO_2_ euthanasia of the animals, the cochleae were extracted, labelled using immunofluorescence and processed for histological analysis. The number of synapses per inner hair cell (IHC) was subsequently quantified using fluorescence microscopy. Furthermore, the serum cortisol levels in all blood samples were determined using a commercial enzyme-linked immunosorbent assay (ELISA).

### 2.3 Electrical stress paradigm

Inescapable electric foot shocks are a well-established stressor in rodent models and have been widely used to induce robust stress responses (Bali & Jaggi, 2015; Santos-Carrasco *et al*., 2025). Prior to each stress session, the animals’ feet were coated with conductive electrode gel. Animals were then placed in a shuttle box consisting of two separate compartments, which were divided by a 6-cm-high partition. The shuttle box was located within an illuminated and sound-attenuated chamber. After a 5-min habituation period, animals assigned to the stress group received electrical foot shocks over a 10-min period, delivered at approximately 60-s intervals with a ±10-s randomized temporal jitter to minimize temporal predictability. Each shock lasted 2 s and was inescapable, as the floors of both compartments of the shuttle box were electrified. During the first session, animals in the stress group received foot shocks for 15 min at approximately 30-s intervals, with added temporal jitter to prevent temporal predictability. Shock intensity was titrated for each animal to ensure reliable perception. Stimulation started at 150 µA and increased in 5 µA increments until a visible behavioural response (e.g. startling or withdrawing) was observed. This intensity was then maintained for the rest of the session. The maximum current did not exceed 230 µA. Animals were continuously monitored via webcam throughout the procedure. Control animals underwent identical handling and chamber exposure but did not receive any electrical stimulation.

As animals in the present protocol were repeatedly exposed to this acute stressor over a period of three weeks, the paradigm is considered to constitute a chronic stress model (Katz *et al*., 1981; Cao *et al*., 2007; Dagyte *et al*., 2009; Atrooz *et al*., 2021). To assess the endocrine stress response, blood samples were collected immediately following selected stress sessions for subsequent cortisol analysis.

### 2.4 Blood sampling and cortisol analysis

Exposure to stressors has been shown to activate the endocrine stress system in Mongolian gerbils, resulting in increased circulating cortisol concentrations (Fenske, 1986; 1996). Blood samples were collected from the femoral vein before the experiment began and within 10 minutes of the first, fifth, tenth and fifteenth stress sessions, as well as before the final GPIAS and ABR measurements were taken. All blood sampling was performed at comparable times around midday to minimise the potential confounding effects of circadian variation in cortisol levels. For blood collection, one experimenter gently restrained the animals while a second experimenter performed femoral vein puncture and sample collection to minimise handling time and stress-related variability. For each sampling time point, only small volumes of blood (around 50–100 µl per animal) were collected in a vial, remaining well within the established animal welfare limits for repeated blood sampling.

The blood samples were then centrifuged at 3000 rpm for 10 minutes and the resulting serum was frozen at -40 °C until further analysis. After completion of the experimental procedures, serum cortisol concentrations were determined using a commercially available ELISA kit (RayBio Human/Mouse/Rat Cortisol Enzyme Immunoassay Kit; RayBiotech Life Inc., USA), following the manufacturer’s instructions. Photometric measurements were performed on a 96-well plate using Multiskan™ FC microplate photometer (Thermo Fisher Scientific, USA) and absorbance values were analysed using SkanIt™ software (Thermo Fisher Scientific, USA).

### 2.5 Acoustic startle response and gap prepulse inhibition

The acoustic startle response (ASR) is a rapid, involuntary motor reflex triggered by sudden, high intensity sounds and controlled by a specific brainstem sensorimotor pathway (Koch, 1999). In rodents, ASR magnitude is quantified as the peak motor reaction to a brief startle stimulus, providing an objective measure of auditory responsiveness that does not require learning or explicit behavioural reports (Clause *et al*., 2011). The ASR can be reliably modulated by preceding sensory cues, forming the basis of prepulse inhibition (PPI) paradigms. In classical PPI, a weak, non-startling auditory stimulus (prepulse) is presented shortly before the startle stimulus (typically 30–300 ms), resulting in a reduction in subsequent ASR amplitude. This phenomenon reflects the presence of intact sensorimotor gating mechanisms, which regulate the processing of incoming sensory information, and has been extensively characterized in rodents and humans (Koch, 1999; Swerdlow *et al*., 2001).

Gap-prepulse inhibition of the acoustic startle reflex (GPIAS) is a variant of the PPI paradigm in which the prepulse is a brief silent gap embedded in continuous background noise. Under normal auditory conditions, detection of this gap suppresses the subsequent startle response. Reduced gap-induced inhibition is widely interpreted as an indicator of tinnitus-like perceptual alterations in rodents, based on the assumption that persistent phantom auditory perception interferes with gap detection (Turner *et al*., 2006; Schilling *et al*., 2017).

Following the experimental protocol, one GPIAS measurement was performed before and after the three-week stress period. Details of the experimental setup and measurement procedures can be found in previous studies (Ahlf *et al*., 2012; Schilling *et al*., 2017; Gerum *et al*., 2019; Tziridis *et al*., 2021). Briefly, the animals were placed in a 15 cm long acrylic tube (inner diameter: 4.3 cm) that was closed at the front with a metal grid and at the rear with a plastic cap. The tube was mounted on a piezoelectric sensor with the front facing the grid, positioned 10 cm from two loudspeakers. The entire setup was located within a sound-attenuated acoustic chamber and the animals were allowed to habituate for 15 minutes prior to testing. GPIAS measurements were performed using narrowband background noise centered at frequencies between 1 – 16 kHz in octave steps to assess frequency-specific gap detection. Startle amplitudes were quantified using standardized procedures and within-subject comparisons between gap and no-gap conditions were used to control for individual differences in baseline startle reactivity. Data processing and GPIAS evaluation were performed using custom-written Python scripts as described in detail by Schilling et al. (2017) and Gerum *et al*. (2019).

### 2.6 Auditory brainstem responses

ABRs were recorded bilaterally in all animals before and after the experimental intervention to assess auditory thresholds and control for potential changes in peripheral auditory sensitivity (Boettcher *et al*., 1993; Tziridis *et al*., 2021). The animals were anaesthetised with a ketamine/xylazine mixture (ketamine 500 mg/kg, xylazine 25 mg/kg), placed in a sound-attenuated chamber and subcutaneous electrodes were positioned at standard recording sites. Acoustic stimuli were presented via a loudspeaker positioned in front of the tested ear. The experimental setup and recording procedures have been described in detail previously (Tziridis *et al*., 2021; Tziridis & Schulze, 2022; Tziridis *et al*., 2024).

### 2.7 Immunohistology and ribbon synapse quantification

After completion of the experiments, animals were euthanized by CO_2_inhalation (Euthanex CO_2_chamber lid, Plexx B.V., Elst, Netherlands) and then decapitated. Both cochleae were rapidly extracted and processed for whole mount immunohistology following established protocols used in gerbil auditory studies (Tziridis *et al*., 2021; Tziridis & Schulze, 2022). Cochlear tissues were fixed, decalcified, and dissected for immunohistochemical analysis. Presynaptic ribbon synapses at IHCs were labelled using antibodies against carboxy-terminal binding protein 2 (CTBP2) (BD Biosciences Cat# 612044, RRID:AB_399431). A Cy3-antibody (Jackson ImmunoResearch Labs Cat# 715-165-150, RRID:AB_2340813) was then applied in accordance with established cochlear synapse quantification procedures (Kujawa & Liberman, 2009). Synaptic structures were then visualized using a fluorescent microscope BZ-9000E (Keyence, Neu-Isenburg, Germany).

Synaptic quantification was performed by counting CTBP2-positive puncta associated with 15 consecutive IHCs along the entire cochlear length by eye (Figure 2). Counts were obtained from both cochleae and synapse counts from the left and right ear were averaged per cochlear region to yield a single synapse count per region per animal. Synapse counts were assigned to characteristic cochlear frequencies based on their distance from the cochlear base, using a Mongolian gerbil–specific cochlear place–frequency map described by Müller (1996). For measuring the distances, we used the BZ Analyzer software from Keyence (Neu-Isenburg, Germany). While the full cochlea was analysed, statistical comparisons focused on regions corresponding to 1, 2, 4, 8, and 16 kHz, matching the frequencies assessed in the GPIAS-measurements.

**Figure 2.**
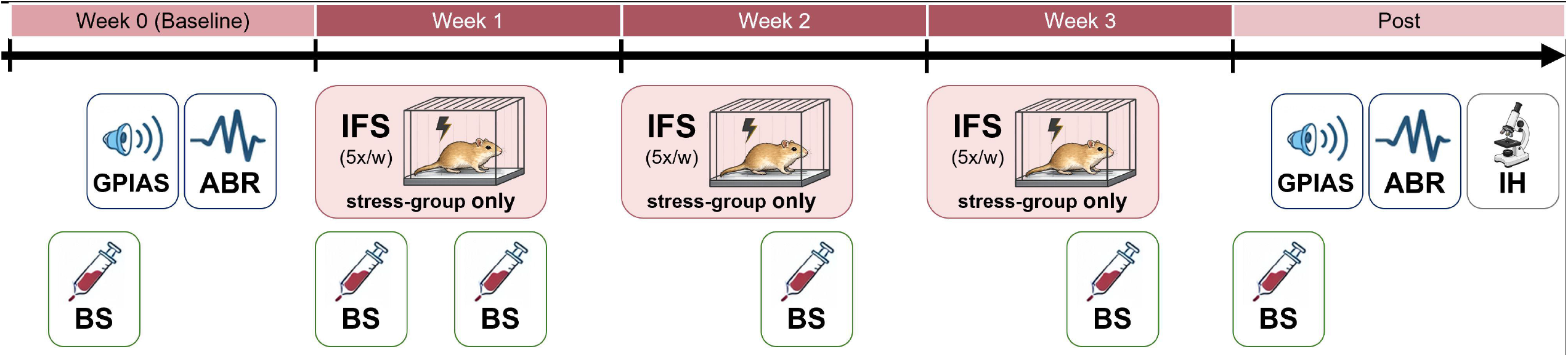
Exemplary microscopic image of cochlear IHC ribbon synapses. Representative microscopic image of cochlear IHCs stained for CTBP2 to visualize ribbon synapses. IHC nuclei are visible and synaptic ribbons appear as discrete puncta beneath the hair cells. Nuclei of outer hair cells (OHCs) are also apparent. For quantification, segments comprising 15 IHCs were analysed and corresponding ribbon synapses were counted by eye. Imaging was performed using a Keyence BZ-9000E microscope with a 40× objective. Scale bar: 50 µm.

### 2.8 Statistical analysis

Statistical analyses were performed using SPSS Statistics (Version 31.0.0.0, IBM Corp.). All tests were two-sided with statistical significance set at p < 0.05.

Changes in serum cortisol concentrations (Δ-cortisol relative to individual baseline) were analysed using a mixed-design ANOVA with time as the within-subject factor and group (stress vs. control) as the between-subject factor. Where appropriate, violations of the sphericity assumption were addressed using Greenhouse–Geisser corrections.

The frequency-specific GPIAS outcomes were evaluated individually first. Animals were subsequently classified as exhibiting tinnitus-like behaviour if a statistically significant reduction in the effect size, defined as the relative suppression of startle amplitude in gap vs. no-gap trials, was observed at any tested frequency relative to baseline (p < 0.05).

Group differences in ABR threshold changes were evaluated using t-tests while frequency-specific effects and potential interactions between group and frequency were assessed using a mixed-design ANOVA with frequency (1, 2, 4, 8 and 16 kHz) as the within-subject factor and group as the between-subject factor.

Cochlear synapse counts were analysed using a linear mixed model with group (stress versus control) and frequency (1, 2, 4, 8 and 16 kHz) as fixed factors. Frequency was modelled as a repeated within-subject factor (subject: animal) using a diagonal covariance structure to allow for frequency-specific variances, while maintaining model parsimony given the moderate sample size. Degrees of freedom were estimated using the Satterthwaite approximation. Estimated marginal means were used for post hoc comparisons.

Exploratory Spearman rank correlation and linear regression analyses were performed to assess potential associations between serum cortisol levels and cochlear synapse counts.

## 3 RESULTS

### 3.1 Cortisol response to repeated stress exposure

Prior to experimental manipulation, baseline serum cortisol concentrations did not differ significantly between stress and control animals (Welch’s t(11.10) = −1.35, p = 0.204), indicating comparable pre-experimental cortisol levels across groups.

A significant main effect of group was detected (between-subjects effect: F(1, 17) = 7.49, p = 0.014), indicating overall differences in Δ-cortisol levels between stress-exposed and control animals averaged across time points.

The interaction between time and group did not reach statistical significance (multivariate test: Pillai’s trace: F(4, 14) = 2.74, p = 0.072). Based on the a priori hypothesis that repeated stress exposure would affect cortisol levels differently over time in stressed animals compared to controls, post hoc pairwise comparisons with Bonferroni correction were conducted to characterise group differences at individual time points further. These analyses revealed significantly higher Δ-cortisol values in stress-exposed animals than in controls on days 5 (p = 0.031) and 10 (p = 0.014). No significant group differences were observed on day 1 (p = 0.079), day 15 (p = 0.104) or at the post-intervention measurement (Figure 3). It can therefore be concluded that while the stress group showed a significant stress response, the control group did not.

**Figure 3.**
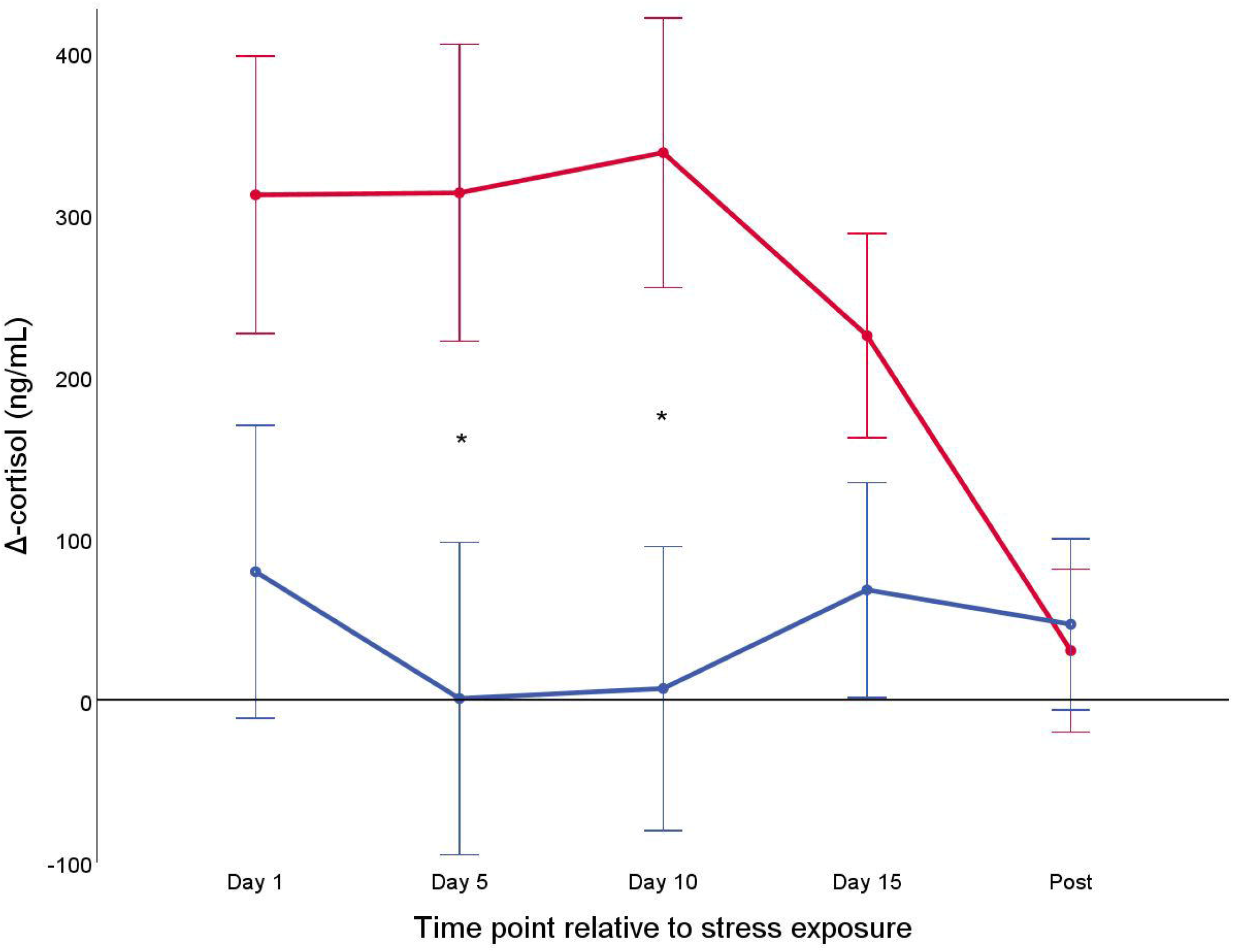
Cortisol response to repeated stress exposure. Changes in serum cortisol concentrations (Δ-cortisol) relative to individual baseline levels are shown for stress-exposed (red) and control (blue) animals at each experimental time point relative to the begin of stress paradigm. Data are shown as mean ± 1 SEM. *Bonferroni-corrected post hoc comparisons revealed significantly higher Δ-cortisol levels in stress-exposed animals compared to controls on Day 5 and Day 10 (p < 0.05).

### 3.2 Behavioural signs of tinnitus

Tinnitus-like behaviour was assessed at five frequencies per animal across all animals, resulting in a total of 100 frequency-specific tests. Only four of these tests (4%) yielded a positive result, highlighting the very low overall prevalence of tinnitus-like responses in the present dataset. The few tinnitus-like classifications were restricted to two discrete frequency bands (1 kHz and 4 kHz), with one animal from each group affected at each frequency, and no animal showing tinnitus-like behaviour across multiple frequencies. As this 4% rate falls within the expected false-positive range at a conventional significance threshold (α = 0.05), we therefore infer that no animal in the present study exhibited reliable tinnitus-like behaviour.

### 3.3 ABR threshold changes

No significant differences in hearing threshold changes were observed between stress-exposed and control animals based on ABR measurements (t(113) = 1.38, p = 0.17). Across the analysed frequency range, neither a frequency-specific effect (F(4, 105) = 0.95, p = 0.44) nor a significant interaction between group and frequency was detected (F(4, 105) = 0.88, p = 0.48). Overall, these findings indicate that repeated stress exposure did not result in measurable alterations in peripheral auditory sensitivity.

### 3.4 Cochlear synapse counts

The analysis revealed no significant main effect of group on synapse counts (F(1, 13.38) = 0.68, p = 0.426), indicating that the number of synapses per inner hair cell was comparable in stress-exposed and control animals. There was also no significant main effect of frequency (F(4, 9.94) = 2.36, p = 0.124) or interaction between group and frequency (F(4, 3.76) = 0.80, p = 0.585), suggesting that synapse counts did not differ systematically across cochlear frequency regions between groups (Figure 4).

**Figure 4.**
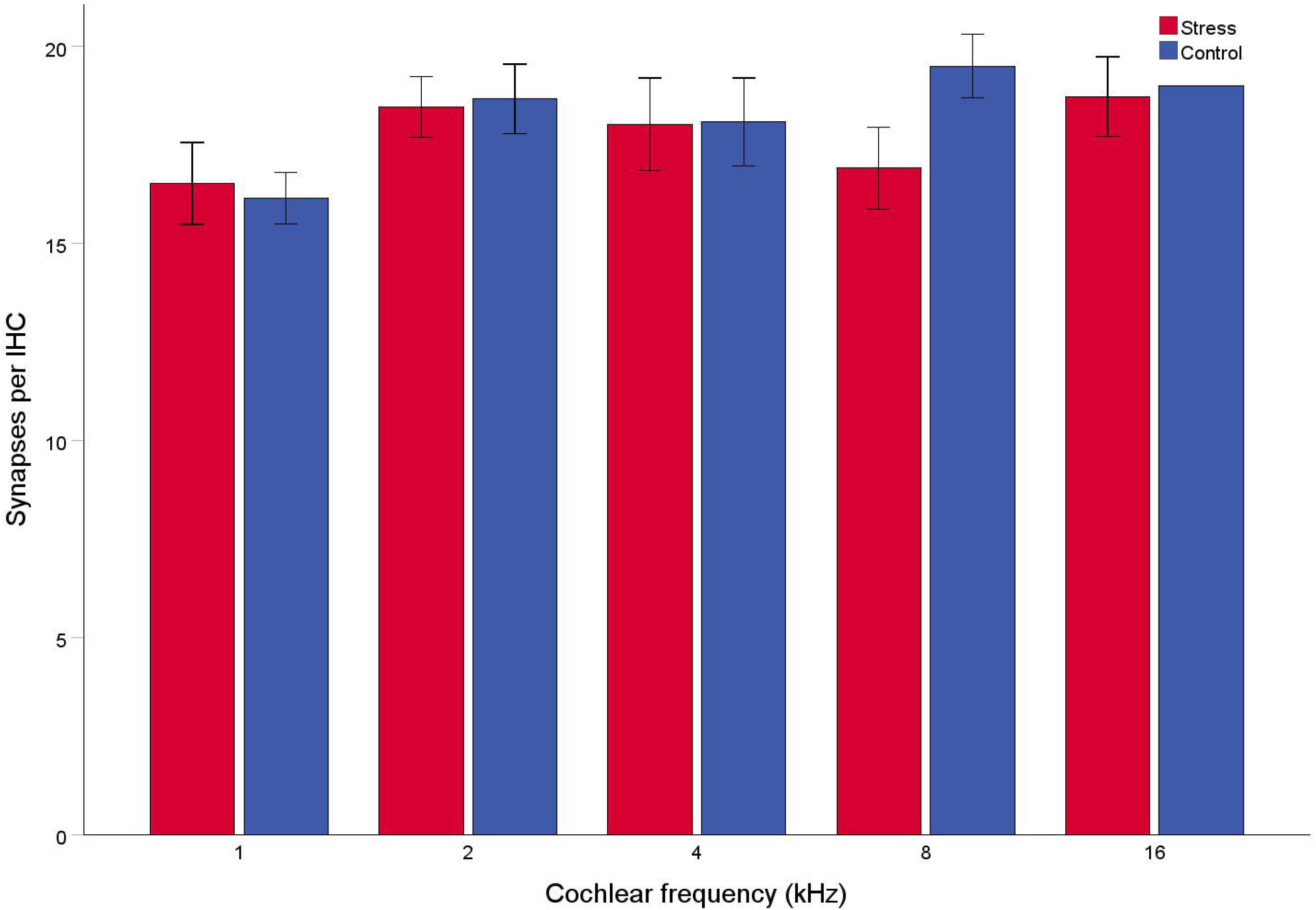
Cochlear ribbon synapse counts across frequency. Mean numbers of ribbon synapses per IHC are shown for stress-exposed (red) and control (blue) animals across cochlear frequency regions (1–16 kHz). Bars represent mean ± 1 SEM. Sample size varied across frequency regions due to preparation-related artefacts. For the control group at 16 kHz, only one animal was available for analysis (n = 1); accordingly, no error bar is shown for this data point.

Consistent with these model-based results, estimated marginal means indicated similar numbers of synapses across the analysed frequency range, with mean synapse counts of 18.28 ± 0.51 in control animals and 17.73 ± 0.44 in stress-exposed animals (mean ± 1 SEM). Pairwise comparisons between groups did not reveal any significant differences (p = 0.426). Taken together, these analyses provide no evidence for stress-induced alterations in cochlear synapse numbers in the present experimental paradigm, although subtle effects of small magnitude cannot be entirely excluded given the moderate sample size.

### 3.5 Further associations

Correlation and linear regression analyses showed no significant associations between serum cortisol concentrations and cochlear synapse counts at any of the experimental time points (day 1, day 5, day 10, day 15, or post-exposure; Spearman r ≤ 0.28, all p > 0.24). No systematic associations were therefore observed between endocrine stress responses and cochlear synapse counts.

## 4 DISCUSSION

We examined whether repeated exposure to stress alone is sufficient to induce tinnitus-like perceptual alterations and cochlear synaptic pathology in a well-established gerbil model. The foot shock paradigm reliably activated the endocrine stress system, as indicated by transient increases in serum cortisol levels immediately after exposure to the shock, which is consistent with the activation of the HPA axis. Given the repeated and inescapable nature of the stressor, the experimental protocol is best interpreted as inducing a state of chronic stress (see Introduction). In line with this, cortisol concentrations in animals exposed to stress returned to control levels after the stress period had ended, which is a well-described pattern in rodent models of repeated homotypic stress. While acute stress reliably elicits transient glucocorticoid elevations, chronic stress is more consistently reflected by sustained alterations in HPA axis regulation and stress responsivity than by persistently elevated basal glucocorticoid levels (Young *et al*., 1990; Kant *et al*., 1992; Rabasa *et al*., 2011; Rostamkhani *et al*., 2012). These alterations in HPA axis function are in line with prolonged adaptation to repeated stress exposure and may indicate a transition toward dysregulation, as described in the later stages of the general adaptation syndrome (Selye, 1936; Herman *et al*., 2016).

Behavioural assessment using GPIAS yielded only rare positive classifications. Across 100 frequency-specific measurements, four met the tinnitus criterion and were evenly distributed between stress and control animals. Furthermore, these detections were confined to isolated frequency bands and did not persist within individual animals. This pattern contrasts with established trauma-based tinnitus models, in which GPIAS deficits are typically robust, frequency-specific and reproducible within subjects (Turner *et al*., 2006; Schilling *et al*., 2017). The observed prevalence (4%) is within the range of false-positive classifications expected at a conventional significance threshold (α = 0.05), indicating that these isolated events are unlikely to reflect consistent tinnitus-like perceptual alterations. While GPIAS is widely applied in rodent tinnitus research, it represents an indirect behavioural proxy and may occasionally yield false-positive outcomes, particularly when event rates are low. This limitation should be considered when interpreting sparse GPIAS effects. Taken together, these findings provide no evidence for the development of reliable tinnitus-like behaviour in any of the animals.

Previous work has reported tinnitus-like GPIAS deficits following exposure to acute and chronic stress (Kim *et al*., 2021; Kim *et al*., 2025). However, in these studies, behavioural testing was performed immediately after stress induction, at a time when acute stress is known to affect startle reactivity and sensorimotor gating (Koch, 1999). For instance, acute stress has been demonstrated to amplify startle responses and diminish prepulse inhibition of the acoustic startle reflex in rodent models, suggesting that stress can directly impact the behavioural indicators employed in GPIAS assays (Santos-Carrasco *et al*., 2025). Similarly, human studies have documented stress-related alterations in sensorimotor gating associated with elevated cortisol levels (Richter *et al*., 2011). These effects may reflect transient changes in attentional or motivational state rather than stable tinnitus-like percepts. In the present study, GPIAS assessments were conducted several days after the final stress session, when serum cortisol levels had returned to baseline. This minimised the likelihood that modulation of behavioural responsiveness by acute stress influenced the outcomes.

Histological analyses converged with the behavioural findings, revealing preserved synaptic integrity across the entire cochlea with no group differences in ribbon synapse counts per IHC. In line with this, ABR measurements provided no evidence of stress-induced hearing impairment. Given that cochlear synaptopathy has been proposed as a key peripheral substrate of tinnitus following auditory injury (Kujawa & Liberman, 2009; Liberman & Kujawa, 2017; Tziridis *et al*., 2021), these results argue against stress-induced structural alterations at the auditory periphery.

These findings are particularly informative considering clinical reports linking stress to the onset or exacerbation of tinnitus. While increased perceived stress and altered cortisol dynamics have been reported in patients (Hébert & Lupien, 2007; Mazurek *et al*., 2012), such observations remain correlational and do not clarify whether stress is a cause or consequence of tinnitus. Here, we directly addressed this question by experimentally isolating stress from auditory trauma and demonstrated that, despite reliable endocrine activation, chronic repeated stress is not sufficient to induce tinnitus-like percepts or cochlear synaptic pathology. From a mechanistic perspective, tinnitus is widely conceptualised as a consequence of maladaptive plasticity following reduced peripheral input (Eggermont & Roberts, 2015). Although stress hormones can modulate neuronal excitability and synaptic transmission (Joëls *et al*., 2011), the present data suggest that these effects do not appear sufficient to initiate tinnitus-related neural reorganization in the absence of auditory injury. Rather than acting as a primary etiological trigger, stress is more likely to influence symptom expression or perceptual salience once auditory dysfunction is present.

Future studies may further disentangle the relationship between stress and tinnitus by experimentally isolating specific stress dimensions or by systematically combining stress exposure with defined auditory vulnerabilities, thereby clarifying the conditions under which stress may modulate tinnitus-related neural plasticity, including potential sex-specific interactions.

## 5 CONCLUSION

In conclusion, the present study shows that repeated exposure to an inescapable foot shock stressor, which constitutes a chronic repeated stress paradigm, is insufficient to induce tinnitus-like perceptual alterations or cochlear synaptic pathology in gerbils when auditory trauma is absent. Although the paradigm reliably activated the endocrine stress system, no behavioural evidence of tinnitus was observed, nor were any structural correlates at the auditory periphery identified. No consistent relationships emerged across endocrine and histological measures. Taken together, these findings argue against a primary causal role of stress alone in tinnitus generation and support the view that tinnitus is more likely to arise from maladaptive neural plasticity following peripheral auditory dysfunction, with stress acting at most as a secondary, modulatory factor.

## ABBREVATIONS

ABR: auditory brainstem response
ASR: acoustic startle reflex
CTBP2: carboxy-terminal binding protein 2
ELISA: enzyme-linked immunosorbent assay
GPIAS: gap-prepulse inhibition of the acoustic startle reflex
HPA: hypothalamic-pituitary-adrenal (axis)
IHCs: inner hair cells
OHCs: outer hair cells
PPI: prepulse inhibition
Δ-cortisol: Change in serum cortisol concentration relative to individual baseline

## Acknowledgements

We are grateful to Dr. Olaf Wendler for the use of lab equipment and to Renate Schäfer for her technical assistance during cortisol measurements. The present work was performed in fulfilment of the requirements for obtaining the degree “Dr. med.” at the Friedrich-Alexander-Universität Erlangen-Nürnberg (FAU).

